# Carbon limitation leads to thermodynamic regulation of aerobic metabolism

**DOI:** 10.1101/2020.01.15.905331

**Authors:** Vanessa A. Garayburu-Caruso, James C. Stegen, Hyun-Seob Song, Lupita Renteria, Jaqueline Wells, Whitney Garcia, Charles T. Resch, Amy Goldman, Rosalie Chu, Jason Toyoda, Emily B. Graham

## Abstract

Organic matter (OM) metabolism in freshwater ecosystems is a critical source of uncertainty in global biogeochemical cycles, yet aquatic OM cycling remains poorly understood. Here, we present the first work to explicitly test OM thermodynamics as a key regulator of aerobic respiration, challenging long-held beliefs that organic carbon and oxygen concentrations are the primary determinants of respiration rates. We pair controlled microcosm experiments with ultrahigh-resolution OM characterization to demonstrate a clear relationship between OM thermodynamic favorability and aerobic respiration under carbon limitation. We also demonstrate a shift in the regulation of aerobic respiration from OM thermodynamics to nitrogen content when carbon is in excess, highlighting a central role for OM thermodynamics in aquatic biogeochemical cycling particularly in carbon-limited ecosystems. Our work therefore illuminates a structural gap in aquatic biogeochemical models and presents a new paradigm in which OM thermodynamics and nitrogen content interactively govern aerobic respiration.

---

Metabolism of organic matter (OM) in freshwater ecosystems plays a large role in global biogeochemical cycles^1–3^, as freshwater ecosystems emit more than 2 Pg C yr^−1^ into the atmosphere^4,5^. These emissions are largely dominated by contributions from river corridors^1,5,6^, and within the river corridor, areas of groundwater-surface water mixing (hyporheic zones) have a disproportionate impact on aerobic respiration^7–9^. Recent field observations have suggested that OM chemistry, and in particular OM thermodynamics, are key to predicting aerobic respiration in hyporheic zones^10–12^. If supported, these observations challenge a widespread paradigm that organic carbon and oxygen concentrations are the primary determinants of aerobic respiration rates and highlight a key source of model uncertainty. Yet, no work has provided direct evidence for OM thermodynamics as a regulator of aerobic respiration in a controlled laboratory environment. Demonstrating this behavior would identify mechanisms that drive field-based phenomena and would enable key properties of OM to be represented in predictive models, thereby contributing to reducing the uncertainty in modeling river corridor biogeochemical cycling^13,14^.

We use highly controlled aerobic microcosms, non-invasive dissolved oxygen consumption rates, and ultrahigh-resolution OM characterization to investigate the role of OM chemistry in determining aerobic respiration in hyporheic zone sediments. Based on field observations^10–12^, we hypothesized that OM chemistry, including thermodynamic favorability and nitrogen (N) content, would regulate aerobic respiration. Historically, investigations of thermodynamic constraints on microbial metabolism have primarily focused on oxidation-reduction reactions that are controlled by the availability of various terminal electron acceptors (e.g., oxygen, nitrate, sulfate)^15–17^. Theory indicates that respiration under aerobic conditions is governed by rate kinetics, while thermodynamic regulation has no influence. This expectation is based on the premise that using oxygen as the terminal electron acceptor provides sufficient energy for ATP generation regardless of thermodynamic properties of the electron donor^18,19^. Consequently, the role of OM thermodynamics in microbial metabolism has been mainly explored under anaerobic conditions^20–23^. However, OM chemistry has recently emerged as a possible regulator of aerobic metabolism based on correlative field-based observation. These studies have suggested that OM thermodynamics interact with N content and carbon concentration to influence aerobic respiration^10–12^. Here, we present the first work to explicitly test OM thermodynamics as a key regulator of aerobic respiration and demonstrate a clear relationship between OM thermodynamic favorability and aerobic respiration under carbon limitation, thus challenging long-held beliefs that respiration rates are govern solely by kinetics.

## Thermodynamic regulation of aerobic respiration

To test the role OM chemistry as a key regulator of aerobic metabolism, we incubated sediments with thermodynamically distinct N-bearing or N-free OM treatments at concentrations commonly observed in freshwater systems (from 0.3 to 9 mg C L^−1^, Supplementary Table 1)^12,24,25^. We inferred carbon limitation at low treatment concentrations (< 3 mg C L^−1^) because sediments were collected from a low carbon area (%C < 0.6 as previously discussed in Graham et al.^11^) with observed C:N < 5^11^ which is typically associated with carbon limitation^26,27^. In addition, sediments were pre-processed prior to incubation until dissolved organic carbon concentrations were below instrument detection (see Methods and Supplementary Fig. 1). Moreover, aerobic respiration increased with treatment concentration from 0.3 to 3 mg C L^−1^ and stabilized between 3 and 9 mg C L^−1^, indicating that additional carbon only stimulated respiration when 3 mg C L^−1^ or less was amended (Fig. 1a, *p* = 0.005 & *p* = 0.43).

**Fig. 1.**
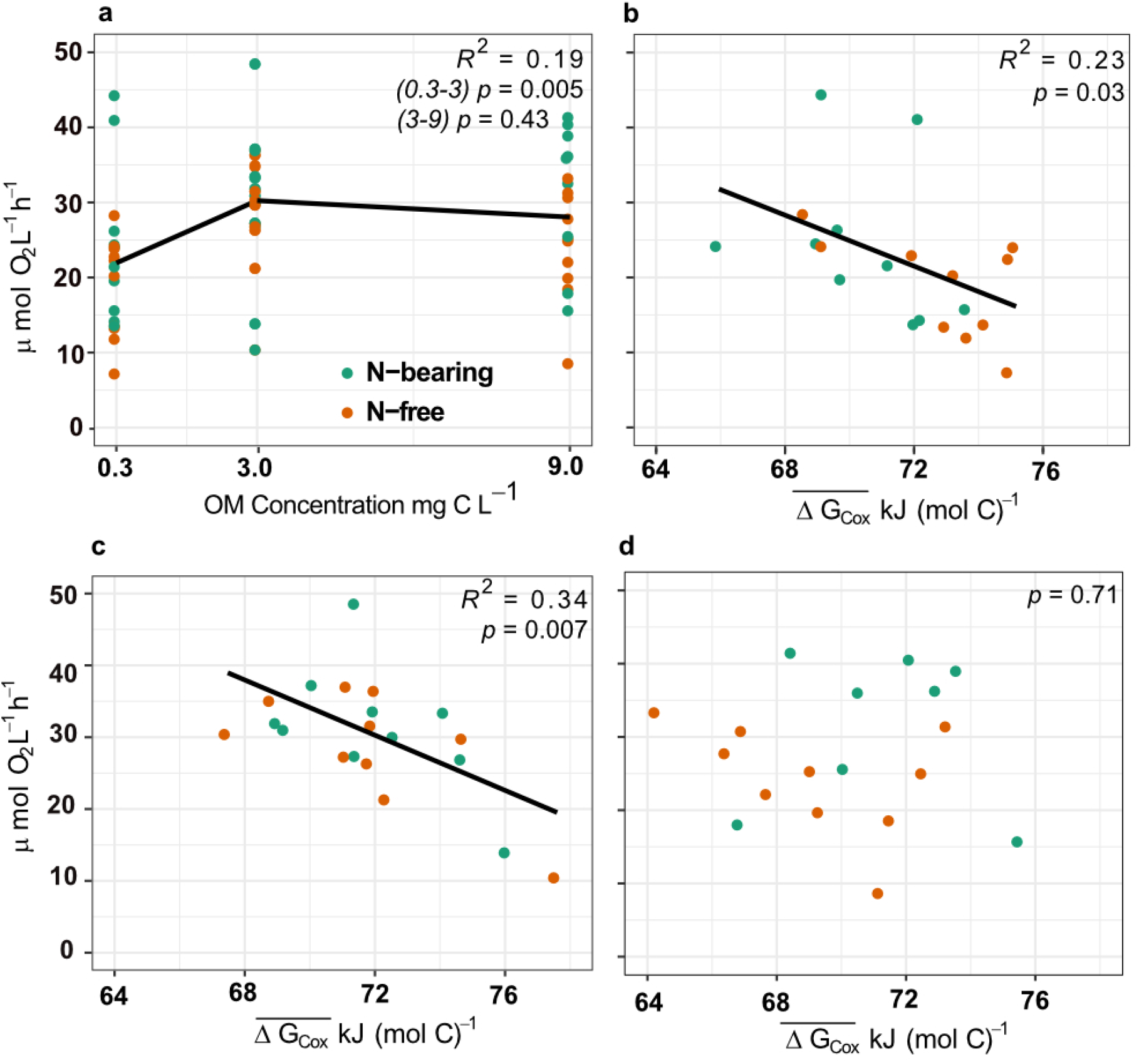
Aerobic respiration rates in microcosms and its relationship with 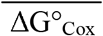. (**a**) Respiration rates were higher in microcosms amended with 3 mg C L^−1^ vs. 0. 3 mg C L^−1^, while microcosms receiving 3 mg C L^−1^ and 9 C mg L^−1^ had similar respiration rates, indicating carbon limitation at low concentrations of amended OM that was alleviated with increasing OM addition. In microcosms with OM added at (**b**) 0.3 C mg L^−1^ and (**c**) 3 mg C L^−1^, 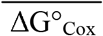 shows a negative relationship with respiration, while microcosms with (**d**) 9 mg C L^−1^ amendments show no relationship between OM thermodynamics and respiration rates.

We provide direct evidence for OM thermodynamics in controlling aerobic respiration by demonstrating that thermodynamically-favorable OM supported enhanced respiration in microcosms under carbon limitation. We use the mean Gibbs free energy of the half reaction of organic carbon oxidation under standard conditions 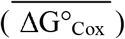 as a proxy for thermodynamic favorability throughout this paper, as per LaRowe and Van Cappellen^28^, Graham et al.^11^, and Stegen et al.^12^. Aerobic respiration was negatively correlated with 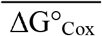 when OM was amended at low concentrations (Fig. 1b-c 0.3 mg C L^−1^ R^2^ = 0.23 *p* = 0.03, 3 mg C L^−1^ R^2^ = 0.34 *p* = 0.007).

While our results support previous field observations that emphasize the role of OM thermodynamics in predicting aerobic respiration^11,12^, we uniquely underscore the central role for OM thermodynamics in the metabolism of carbon-limited ecosystems using highly controlled laboratory experiments. A field study by Graham et al.^11^ showed that thermodynamically favorable OM was preferentially metabolized in sediments across a vegetation gradient. Similarly, Stegen et al.^12^ highlighted the importance of OM thermodynamics in regulating aerobic respiration within hyporheic zones. While the previous works are based on correlative observations, our controlled laboratory microcosms allowed us to clearly demonstrate thermodynamic regulation of aerobic metabolism and define the conditions under which OM thermodynamics provide an avenue for improving aquatic biogeochemical models.

Specifically, we reveal a dependency of thermodynamic OM regulation on organic carbon concentration, as sediments putatively not experiencing carbon limitation had no evidence of thermodynamic constraints on aerobic respiration (Fig. 1d, 9 mg C L^−1^ *p* = 0.71). Low molecular weight carbon compounds, such as the amino and organic acids used for treatments in this experiment (see Methods), are highly bioavailable to microorganisms which have direct uptake pathways for these molecules^29–31^ in contrast to polymeric OM in sediments that requires the production of extracellular enzymes^32,33^. Thus, we hypothesize that excess low molecular weight and highly bioavailable OM diminishes the benefits of thermodynamic preference in metabolism, because the low microbial cost of direct uptake outweighs the energy benefit gained from selective carbon metabolism.

## Pathways of OM metabolism vary with thermodynamic control of aerobic respiration

We underscore a need for improved model structures for predicting aerobic respiration, as we observed different metabolic processes between carbon-limited and carbon-replete environments. To investigate metabolic processes involved in aerobic respiration, we calculated inferred biochemical transformations in ultrahigh-resolution OM profiles following procedures described previously^10–12,34–36^. This method relies on the mass accuracy of Fourier transform ion cyclotron resonance mass spectrometry (FTICR-MS) and produces a count of the number of times a specific molecule (e.g., glucose, valine, glutamine, etc.) is putatively gained or lost in reactions (see Methods).

While previous work has suggested both thermodynamic and N-related regulation of aerobic respiration^10–12^, we postulate a carbon limitation threshold beyond which thermodynamic controls do not persist. Our data suggest that beyond this threshold (i.e., in the absence of carbon limitation), aerobic respiration is coupled to organic N metabolism. We found no differences in OM chemistry (Supplementary Fig. 2a-b, all *p* > 0.05), respiration rates (*p* = 0.11 at 0.3 mg C L^−1^ and *p* = 0.24 at 3 mg C L^−1^) or biochemical transformations (Fig. 2a-b, 0.3 mg C L^−1^ *p* = 0.67, 3 mg C L^−1^ *p* = 0.91) across microcosms with N-bearing vs. N-free OM amended when respiration was thermodynamically-controlled (i.e., in carbon limited conditions). However, respiration in carbon-replete microcosms (9 mg C L^−1^) increased with the addition of organic N relative to N-free OM, and pathways of OM metabolism varied between treatments with or without added organic N (Fig. 2c-d, *p* = 0.04, *p* = 0.04). Biochemical transformations in microcosms receiving 9 mg C L^−1^ more frequently involved N when organic N was added to microcosms (Supplementary Fig. 3, *p* = 0.02). Additionally, we provide evidence that N-enriched molecules are preferably consumed in natural environments with surplus carbon. Bioavailable organic N addition increased the relative abundance of more complex protein-like OM, suggesting the preservation of sediment-bound OM containing N in the presence of more accessible N sources (Supplementary Fig. 2c, *p* < 0.01). Together, these results are consistent with N-mining observed previously in this system, whereby OM is oxidized for microbial acquisition of N^10,37^.

**Fig. 2.**
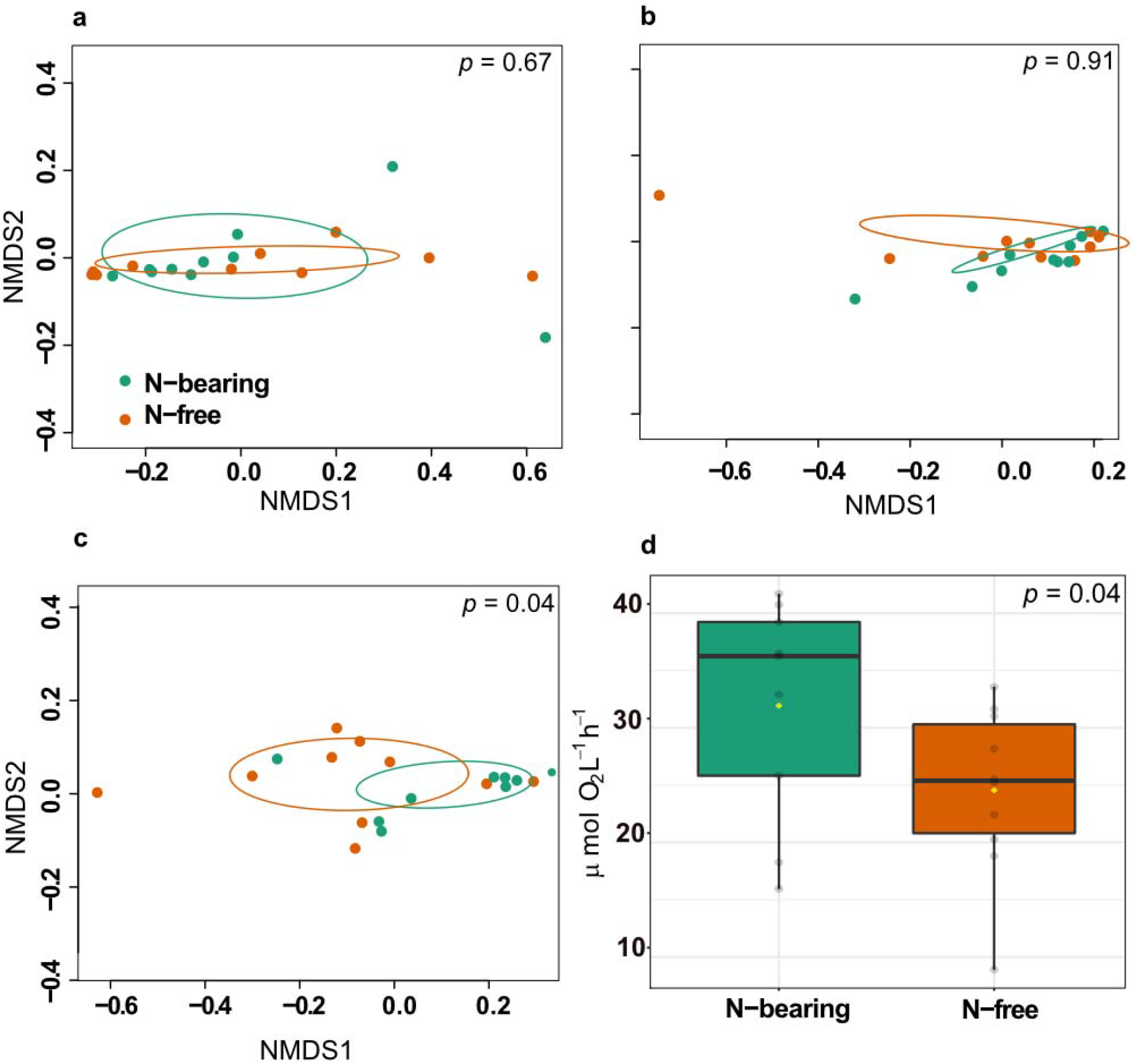
Biochemical transformations between N-bearing and N-free OM amendments and its link with respiration rates. Non-metric multidimensional scaling (NMDS) plots of microcosms receiving low OM concentrations (**a**) 0.3 mg C L^−1^ and (**b**) 3 mg C L^−1^ showed no difference in biochemical transformation profiles between N-bearing and N-free amendments. In contrast, microcosms receiving high OM concentrations (**c**) 9 mg C L^−1^ had significantly different biochemical transformation profiles between N-bearing and N-free amendments. (**d**) Microcosms amended with N-bearing OM at 9 mg C L^−1^ also showed enhanced respiration vs. those receiving N-free OM at the same concentration. In contrary, respiration rates were not statistically different between N-bearing and N-free microcosms receiving low OM (0.3 and 3 mg C L^−1^). Colors in all panels indicate N-bearing (teal) and N-free (orange) OM amendments. *P*-values in (**a-c**) were derived from PERMANOVAs, and the *p*-value in (**d**) was calculated using a one-sided Mann-Whitney U test.

More broadly, we highlight that when organic carbon is in excess, there is a dependency of aerobic respiration on specific nutrient limitations rather than on OM thermodynamics. Therefore, OM thermodynamics appear to be most informative of aquatic biogeochemistry within carbon-limited ecosystems, while N availability governs aerobic respiration at higher carbon to nitrogen ratios (C:N). While N-dependent aerobic respiration at high C:N is consistent with nutrient limitations observed in variety of systems^38,39^, thermodynamic regulation at low C:N ratios challenges the widespread notion that organic carbon and oxygen concentrations are the main variables driving respiration.

## Enhancing model predictions with a new paradigm of aquatic biogeochemical cycling

While many processed-based models represent biogeochemical cycles, most model structures contain only a few lumped carbon pools that do not fully represent the complexity of natural OM sources^20,21^. Organic matter cycling is typically modeled following Michaelis-Menten kinetics, with OM separated into particulate or dissolved pools^40^. In some cases, these pools are further categorized by environmental properties or bioavailability, and each subpool is assigned a fixed mineralization rate^20,40–42^. These traditional approaches do not address OM chemistry because commonly used bulk characterization techniques do not provide sufficient molecular detail, and because representing individual OM molecules in a given ecosystem is computationally unfeasible. Our work demonstrates that processes associated with OM thermodynamics and N content strongly influence aerobic respiration—and, in turn, biogeochemical reaction networks— thus providing a link between OM chemistry and biogeochemical rates that will help inform and parameterize new and existing models.

We therefore propose a new conceptual model with direct avenues for model incorporation in which OM thermodynamics regulates aerobic respiration until organic carbon concentrations are sufficient to induce nutrient limitations (Fig. 3). We suggest that a combination of organic carbon concentration and thermodynamic limitation governs aerobic respiration in sediments with low carbon to nutrient ratios. In contrast, nutrient limitation regulates respiration when bioavailable carbon is in excess, and in this scenario, organic N may be a key constraint on respiration. This is consistent with previous reports of a strong role from organic N cycling in hyporheic zones and freshwater systems^10,12,43,44^. While our work represents a single system, our conceptual model is meant to provide proof-of-concept for more spatially extensive studies that will allow broader transferability.

**Fig. 3.**
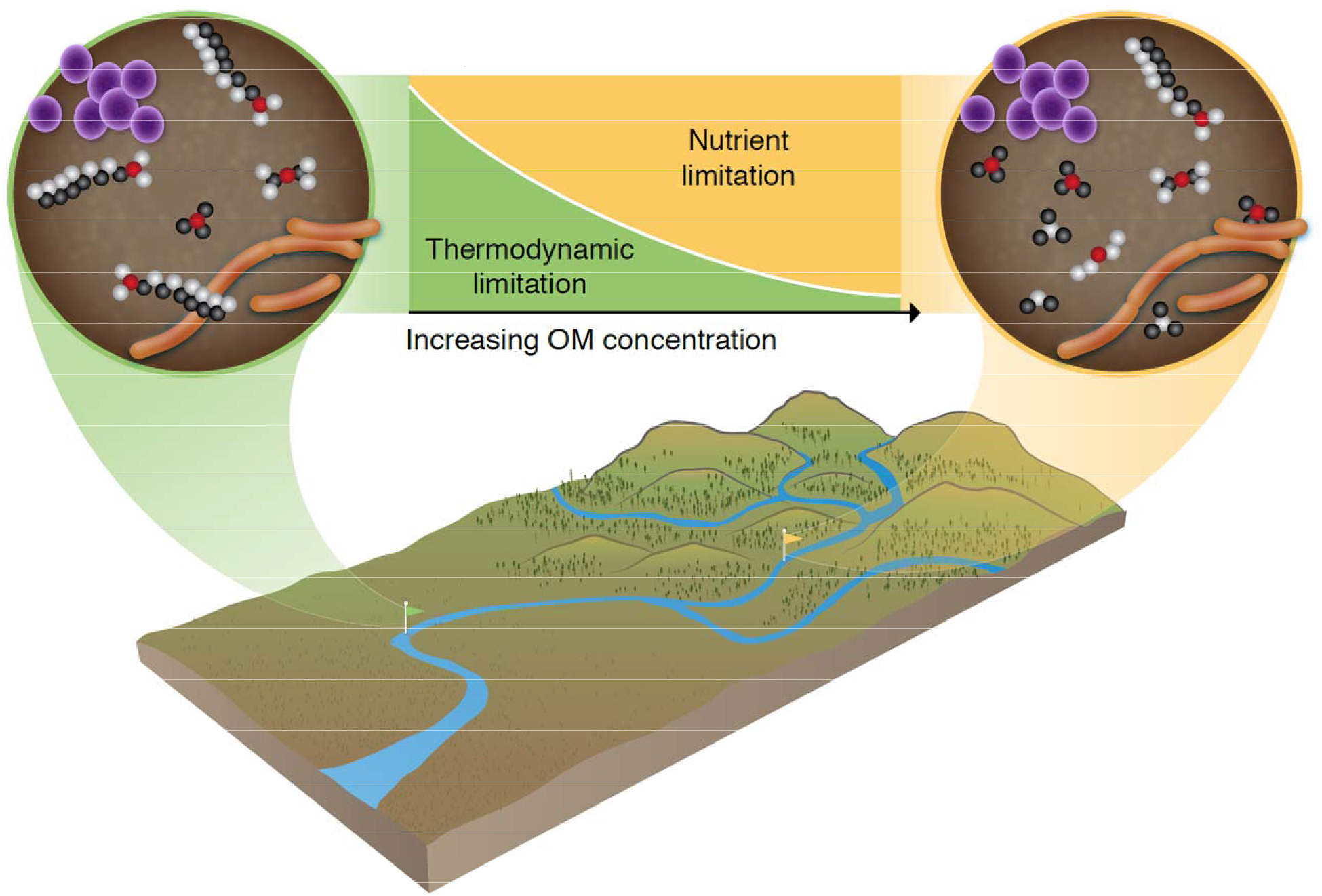
Conceptualization of thermodynamic and nutrient regulations on aerobic respiration. We propose a new conceptual model in which thermodynamic and nutrient limitations dually control aerobic respirations. We suggest that thermodynamic properties of OM govern aerobic respiration rates in ecosystems with low carbon to nutrient ratios. When OM concentration reaches a threshold, thermodynamic controls do not persist, and nutrient availability, particularly N regulate respiration. This work highlights a structural gap in aquatic biogeochemical models and challenges long-held beliefs about aerobic metabolism being solely governed by reaction kinetics. This new paradigm provides a link between OM chemistry and biogeochemical rates with direct avenues for model incorporation, where OM chemistry regulates OM oxidation through its thermodynamic properties until OM concentrations are sufficient to induce nutrient limitations.

To our knowledge, this is the first work to provide direct evidence for OM thermodynamics as a regulator of aerobic respiration in a controlled laboratory environment and highlights a key gap in current mechanistic understanding of OM cycling. In order to improve model accuracy, we reveal a need to explicitly represent the interactions between OM thermodynamics, nutrient limitations, and organic carbon concentration in process-based models of aquatic carbon cycling. Our results constrain the environments under which specific chemical attributes of OM are valuable for rate predictions and provide guidance on the data types that are needed to accurately represent hyporheic zone biogeochemistry. These processes cannot be represented by lumped OM pools informed by decades of coarse measurements, and thus we highlight the utility of high-resolution technologies in models of aquatic biogeochemistry and present a new paradigm in which aerobic respiration is governed by a combination of carbon concentration, OM thermodynamics, and nutrient limitations.

## Supporting information

Supplementary Table 2. List of transformations

## Acknowledgements

This research was supported by the U.S. Department of Energy (DOE), Office of Biological and Environmental Research (BER), as part of Subsurface Biogeochemical Research Program’s Scientific Focus Area (SFA) at the Pacific Northwest National Laboratory (PNNL). Data were generated under EMSL user proposal 51180. A portion of the research was performed at Environmental Molecular Science Laboratory User Facility. PNNL is operated for DOE by Battelle under contract DE-AC06-76RLO 1830.

## Author Contributions

V.G.C., E.B.G, H.S.S. and J.C.S., conceptualized the study; V.G.C., L.R., J.W., W.G, and A.G., carried out the study; C.T.R., R.C. and J.T. conducted instrumental analyses; V.G.C. and E.B.G drafted the manuscript and all authors contributed to the writing.

## Materials and methods

### Study site and sediment collection

This study was conducted using sediments from the Columbia River hyporheic zone within the Hanford Site 300 Area (approximately 46° 22’ 15.80”N, 119° 85 16’ 31.52”W) in eastern Washington, USA^10,11,45^. Hyporheic zone sediments were collected in April 2018 at five locations, separated by ~2 m (depth: ~30 cm). Sediments were sieved in the field to < 2 mm, homogenized, and kept on ice until same-day laboratory processing. We performed sequential organic carbon extractions with synthetic river water (see Supplementary Methods for composition) prior to incubations to minimize the influence of carbon mobilized from sediments during field sampling. Details regarding pre-processing step are provided in the Supplementary Methods.

### Laboratory Microcosms

We used a total of 85 microcosms in a full factorial design (a) four chemically distinct OM amendments at three concentrations, (b) four autoclaved controls (heat kills), and (c) one synthetic water control, each treatment with five replicates. Incubations were performed over the course of 5 days, where each day we incubated 17 bioreactors (i.e., 1 replicate treatment per day, see design in Supplementary Table 1). On the day prior to the experiment, 10 g of pre-processed sediments were removed from 4°C storage and subsampled into 20 mL borosilicate glass vials. Vials were left in the dark at ambient laboratory temperature for 8 h before incubation to acclimate to room temperature.

To initiate microcosms, 18 mL of treatment solution was added to vials containing sediment, leaving <1 mL headspace. Treatment solution consisted of synthetic river water and the specific OM compound at the desired concentration (Supplementary Table 1). Nitrate and phosphate were added to the synthetic river water to provide sufficient nutrients for the duration of the experiment - nitrate concentration matched ambient groundwater while phosphate concentration matched the Redfield ratio relative to groundwater N (16N:P). We added the following 4 types of OM because their thermodynamic properties encompassed the extremes experienced by the surface and the groundwater in situ^12^ and were either N-bearing or N-free: Lysine (Gibbs free energy of the half reaction of organic carbon under standard conditions, ΔG°_cox_ = 79.40 kJ (mol C)^−1^, N-bearing), Serine (ΔG°_Cox_ = 41.21 kJ (mol C)^−1^, N-bearing), Propionate (ΔG°_Cox_=79.40 kJ (mol C)^−1^, N-free), and Ascorbate (ΔG°_Cox_ = 41.21 kJ (mol C)^−1^, N-free). The vials were placed horizontally on a shaker at 250 rpm in the dark at 21±1 °C for the duration of the experiment, except during dissolved oxygen measurements.

### Respiration rates

Dissolved oxygen (DO) concentration (μmol L^−1^) was measured in each microcosm every hour for 6 h using 0.5 cm diameter factory-calibrated oxygen sensors and an oxygen optical meter (Fibox 3; PreSens GmbH, Regensburg, Germany). The DO measurements were automatically corrected for temperature and the data were recorded using PST3v602 software (PreSens GmbH). Respiration rates were calculated as the slope of the linear regression between DO concentration and incubation time for each microcosm (Supplementary Fig. 4-6). We infer that changes in DO were driven by aerobic respiration, as DO in heat kills did not change during the incubation (Supplementary Fig. 7). pH measurements were also collected using an optical meter and factory calibrated pH sensor spots (pH-1 mini; PreSens GmbH). pH values did not change during the incubation (Supplementary Fig. 8). After 6 hours, microcosm contents were transferred to 50 mL sterile polypropylene centrifuge tubes and centrifuged for 5 min at 3200 rcf and 20°C. The supernatant was filtered through a 0.22 μm polyethersulfone membrane filter (Millipore Sterivex, USA) frozen at −20°C until further analysis.

### Fourier transform ion cyclotron resonance mass spectrometry (FTICR-MS)

Fourier transform-ion cyclotron resonance mass spectrometer (FTICR-MS) (12 Tesla (12T) Bruker SolariX, Billerica, MA) located at the Environmental Molecular Sciences Laboratory in Richland, WA, was used to collect high-resolution mass spectra of the OM. Resolution was 220K at 481.185 m/z. The FTICR-MS was outfitted with a standard electrospray ionization (ESI) source, and data was acquired in negative mode with the voltage set to +4.4kV. Data were collected with an ion accumulation time of 0.3 sec from 98 – 900 m/z at 4M. One hundred fourty-four scans were co-added. BrukerDaltonik (version 4.2) was used to convert raw spectra to a list of m/z values by applying FTMS peak picker module. Chemical formulas were then assigned using in-house software following the Compound Identification Algorithm^46–49^, using the criteria previously described by Graham et al.^10,11^. The chemical character of the compounds identified in the FTICR-MS spectrum and their biochemical classes were evaluated using Van Krevelen diagrams.

We calculated the ΔG°_Cox_ to evaluate relationships between aerobic respiration and OM thermodynamics, as per LaRowe and Van Cappellen^28^. An expanded description of sample preparation, instrument and FTCR-MS data processing for estimating Van Krevelen diagrams and ΔG°_Cox_ is presented in the Supplementary Methods.

### Identification of biochemical transformations using FTICR-MS

Biochemical transformations were inferred by calculating all possible pairwise mass differences within a sample’s spectrum and matching differences (within 1□ppm) to a list of common biochemical transformations^50^. Biochemical transformations were identified following the procedures described by Breitling et al.^50^ and previously employed by Bailey et al.^34^, Graham et al.^10,11^, Moritz et al.^36^, Kaling et al.^35^, and Stegen et al.^12^. Briefly, pairwise mass differences between all m/z peaks in a sample were compared with a reference list of 1298 commonly observed biochemical reactions of organic matter (Supplementary Table 2). For mass differences matching to compounds in the reference list, we inferred the gain or loss of that compound via a biochemical transformation.

### Statistical analyses

All statistical analyses were completed using R (version 3.4.1). Linear regressions were used to assess the relationship between respiration rates and 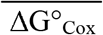. Differences across groups were evaluated with one-sided Mann-Whitney U test. Permutational multivariate analysis of variance (PERMANOVA) of Bray–Curtis distances was used to assess dissimilarities among biogeochemical transformations in the “vegan” R package. PERMANOVAs were stratified by ΔG°_Cox_ to account for any differences due to the thermodynamic properties of the treatment solution. Biochemical transformations were visualized with non-metric multidimensional scaling (NMDS).

