## Supplementary Information for "Carbon limitation leads to thermodynamic regulation of aerobic metabolism"

**Supplementary Methods**

**Sediment collection and processing**

This study was conducted using low carbon and low nitrogen (N) sediments^1^ from the Columbia River hyporheic zone within the Hanford Site 300 Area in eastern Washington, USA. In order to minimize the influence of carbon mobilized from sediment sampling, we performed sequential organic carbon extractions with synthetic river water prior to incubations. The concentration of ions in the synthetic river water solution (mmol L^−1^) was 1.8 Na^+^, 0.4 Ca^2+^, 0.2 Mg^2+,^ 0.02 K^+^, 1.6 HCO_3_^−^, 1.4 Cl^−^, 0.02 NO_3_^−^, and 0.1 SO_4_^2-^. 35 g of sediments were subsampled into 500 mL polypropylene centrifuge bottles and 350 mL of synthetic river water were added. The bottles were continuously shaken in the dark at 250 rpm and 21°C for 1 h, after which the bottles were centrifuged at 6580 rpm and 21°C for 5 min. The supernatant volume was exchanged with new synthetic river water and the bottles were placed again on the shaker to begin a new extraction. On average, we exchanged the volume of the bottles seven times a day for a total of 21 extractions. The supernatant was collected at three time points per day and filtered through 0.22 μm polyethersulfone membrane filter (Millipore Sterivex, USA) into borosilicate glass vials, for dissolved non-purgeable organic carbon analysis (NPOC; Shimadzu combustion carbon analyzer TOC–Vcsh with ASI– V auto sampler). Extractions were performed for three days, until NPOC measurements in the supernatant was near instrument detection limit (0.3 mg C L^-1^, Supplementary Fig. 1). After the extractions, the sediments were either subsampled for following day’s experiment or stored in dark at 4°C for 24 h to be used on the next day’s incubation.

**Laboratory incubations**

Supplementary Table 1 displays the full factorial design of microcosm incubations and treatments with thermodynamically distinct N-bearing or N-free treatment solution at concentrations commonly observed in freshwater systems^2-4^. Nitrate and phosphate were added to the synthetic river water to provide sufficient nutrients for the duration of the experiment, the final concentration of ions in the solution used for the microcosms (mmol L^−1^) was 1.8 Na^+^, 0.4 Ca^2+^, 0.2 Mg^2+,^ 0.02 K^+^, 1.6 HCO_3_^−^, 1.4 Cl^−^, 0.34 NO_3_^−^, 0.03 PO_4_^−^ and 0.1 SO_4_^2-^. Sediment heat kills were prepared the day before the experiment by autoclaving 40 g of pre-treated sediments for 90 min in gravity cycle at 121 ºC and 15 psi. All incubations were performed in 20 mL borosilicate glass vials certified to meet the EPA’s volatile organic analysis standards.

**Fourier transform ion cyclotron resonance mass spectrometry (FTICR-MS)**

Supernatant collected after incubation was acidified to pH 2 with 85% phosphoric acid and extracted with PPL cartridges (Bond Elut), following Dittmar et al.^5^. Subsequently, a 12 Tesla (12T) Bruker SolariX FTICR-MS (Bruker, SolariX, Billerica, MA) located at the Environmental Molecular Sciences Laboratory in Richland, WA, was used to collect high-resolution mass spectra of the OM.

Van Krevelen diagrams were used to visualize and compare the average properties of organic compounds and assign compounds based on elemental composition to major biochemical classes (e.g., lipid-, protein-, lignin-, carbohydrate-, and condensed aromatic-like). Compounds were plotted as a function of their molar H:C ratios (y-axis) and molar O:C ratios (x-axis)^6^. We used the Van Krevelen biochemical compound classes to estimate the relative abundance of each metabolite class in a sample.

We calculated the Gibbs free energy of the half reaction of organic carbon oxidation under standard conditions (ΔG^º^_Cox_) to evaluate relationships between aerobic respiration and OM thermodynamics. Although ΔG^º^_Cox_ varies with the availability of the electron acceptor, our experimental design maintained homogenous aerobic conditions in each microcosm using continuous shaking, such that oxygen was putatively the terminal electron acceptor, allowing us to make direct comparisons across samples using only ΔG^º^_Cox_.

As per LaRowe and Van Cappellen^7^, we first estimated the nominal oxidation state of carbon (NOSC) with the following equation:

$$NOSC=-\left( \frac{-Z+4a+b-3c-2d+5e-2f}{a} \right)+4 (1)$$

Where, $Z$ is the net charge and $a$, $b$, $c$,$d$, $e$, and $f$ are the number of atoms of elements C, H, N, O, P, and S, respectively, in a given organic compound. Subsequently, ΔG^º^_Cox_ was estimated from the empirical equation:

$$\Delta G_{Cox}^{o}=60.3-28.5\left( NOSC \right) (2)$$

ΔG^º^_Cox_ values are usually positive, indicating that the half reaction of organic carbon oxidation must be coupled to the half reaction of reduction of a terminal electron acceptor. ΔGº_Cox_ values are more thermodynamically-favorable when they are closer to zero. Mean ΔG^º^_Cox_ of all OM species present in a sample’s FTICR-MS spectra (i.e., all unique m/z peaks) was calculated and used in downstream analyses.

**Supplementary Tables**

**Supplementary Table 1** Factorial design of treatments for incubations.

|  | Control | N-bearing compounds | | N-free compounds | |
| --- | --- | --- | --- | --- | --- |
|  | *Synthetic water* | *Lysine* | *Serine* | *Propionate* | *Ascorbate* |
| Amended OM concentration  (mg C Lˉ¹) | 0 | 0.3 | 0.3 | 0.3 | 0.3 |
|  |  | 3 | 3 | 3 | 3 |
|  |  | 9 | 9 | 9 | 9 |
| Heat kill OM concentration  (mg C Lˉ¹) |  | 9 | 9 | 9 | 9 |
| ΔG^º^_Cox_ kJ (mol C ^-1^) |  | 79.40 | 41.21 | 79.40 | 41.21 |

**Supplementary Table 2** List of Biochemical transformations.

File name: Supplementary Table 2_List of transformations.xlsx

Description: List of biochemical transformation (i.e., masses gained or lost) that can be inferred from FTICR-MS data.

**Supplementary Figures**

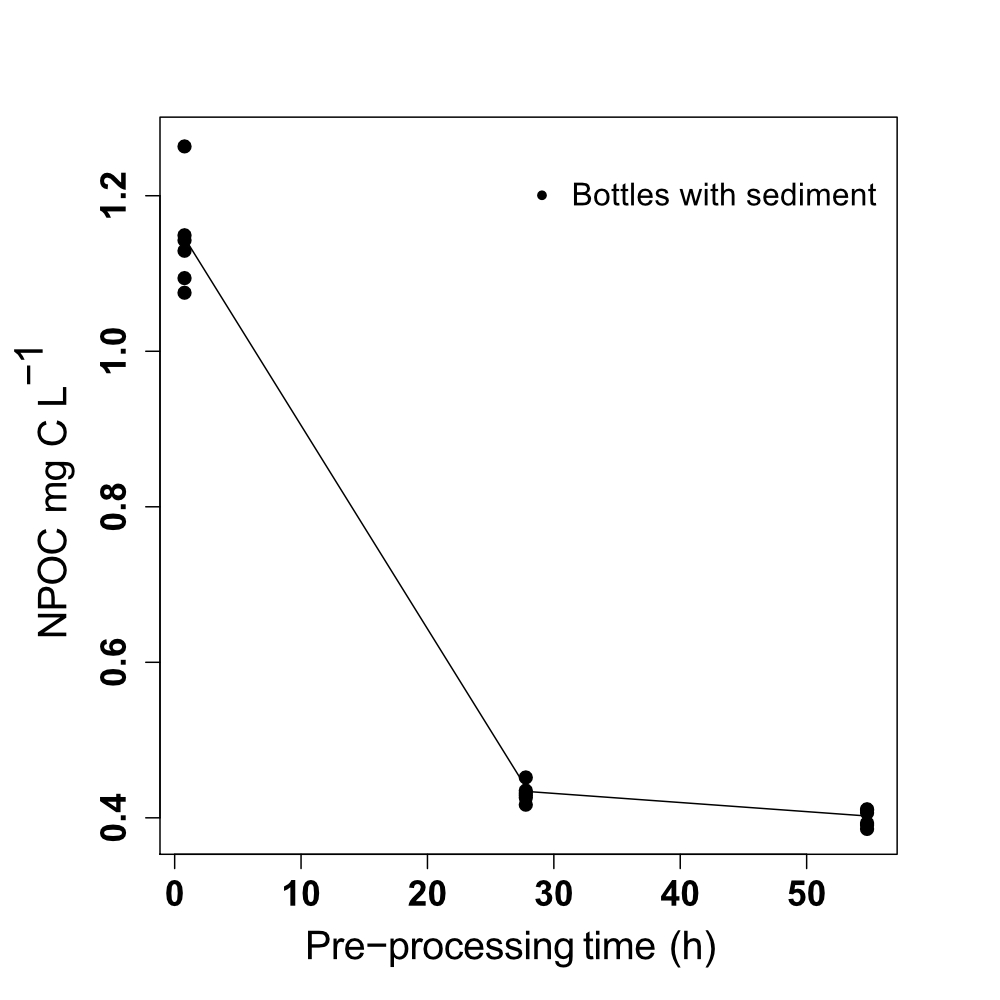

**Supplementary Fig. 1** Supernatant non-purgeable organic carbon (NPOC) concentration from sediment-synthetic water extractions. Extractions were performed for three days prior to incubation, until NPOC measurements in the extracted supernatant was near instrument detection limit (0.3 mg C L^-1^).

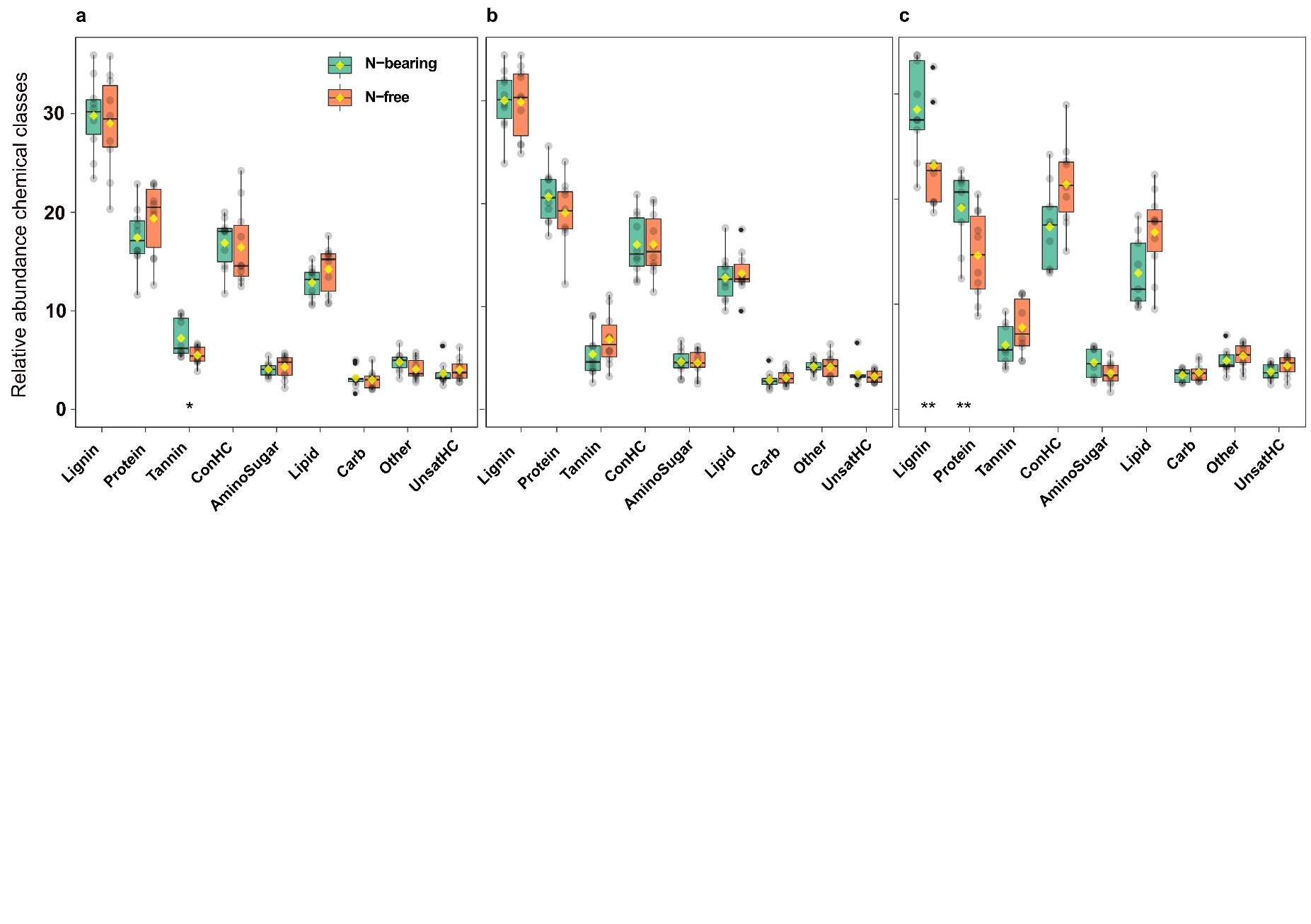

**Supplementary Fig. 2** Relative abundance of Van Krevelen chemical classes in N-bearing and N-free OM amendments. Treatments with low OM concentration, (**a**) 0.3 mg Lˉ¹ C and (**b**) 3 mg Lˉ¹ C, show no statistical difference between metabolite groups identified in each sample. (**c**) 9 mg Lˉ¹ C, shows increased abundance of protein-like compounds in N-bearing relative to N-free microcosms, suggesting potential N-mining. Colors indicate different OM amendments (N-free vs N-bearing). Plots are marked with a center black line in each box (median), yellow dot (mean), and hinges (values at 25^th^ and 75^th^ percentiles). *P*-value was derived from one-sided Mann-Whitney U test. One asterisk (*) indicates *p* ≤ 0.05, and two (**) indicates *p* ≤ 0.01.

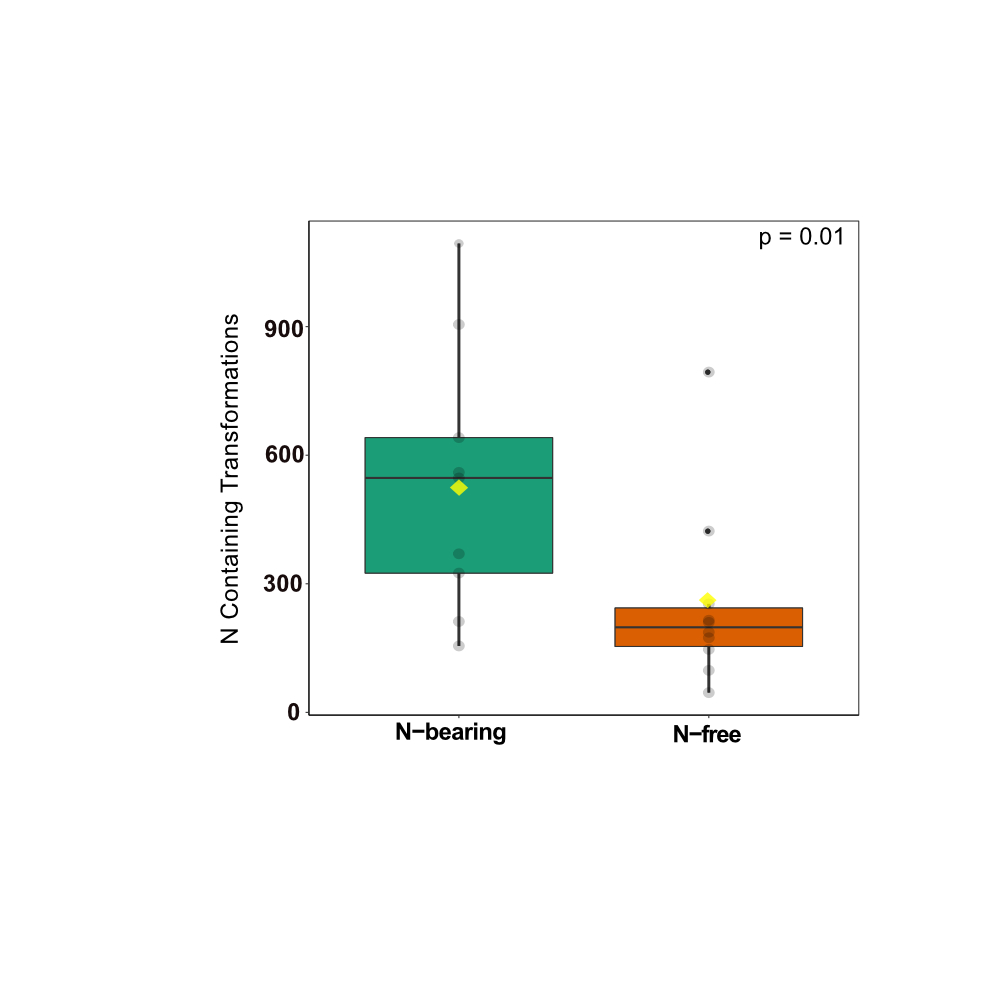

**Supplementary Fig. 3** Number of biochemical transformations involving gain or loss of a N containing molecule**.** N-bearing treatments exhibited a higher number of biochemical transformations involving N than the N-free treatments, emphasizing the relevance of the N cycle in carbon replete microcosms. Colors indicate different OM amendments (N-free vs N-bearing). Plots are marked with a center black line in each box (median), yellow dot (mean), and hinges (values at 25^th^ and 75^th^ percentiles). *P*-value was derived from one-sided Mann-Whitney U.

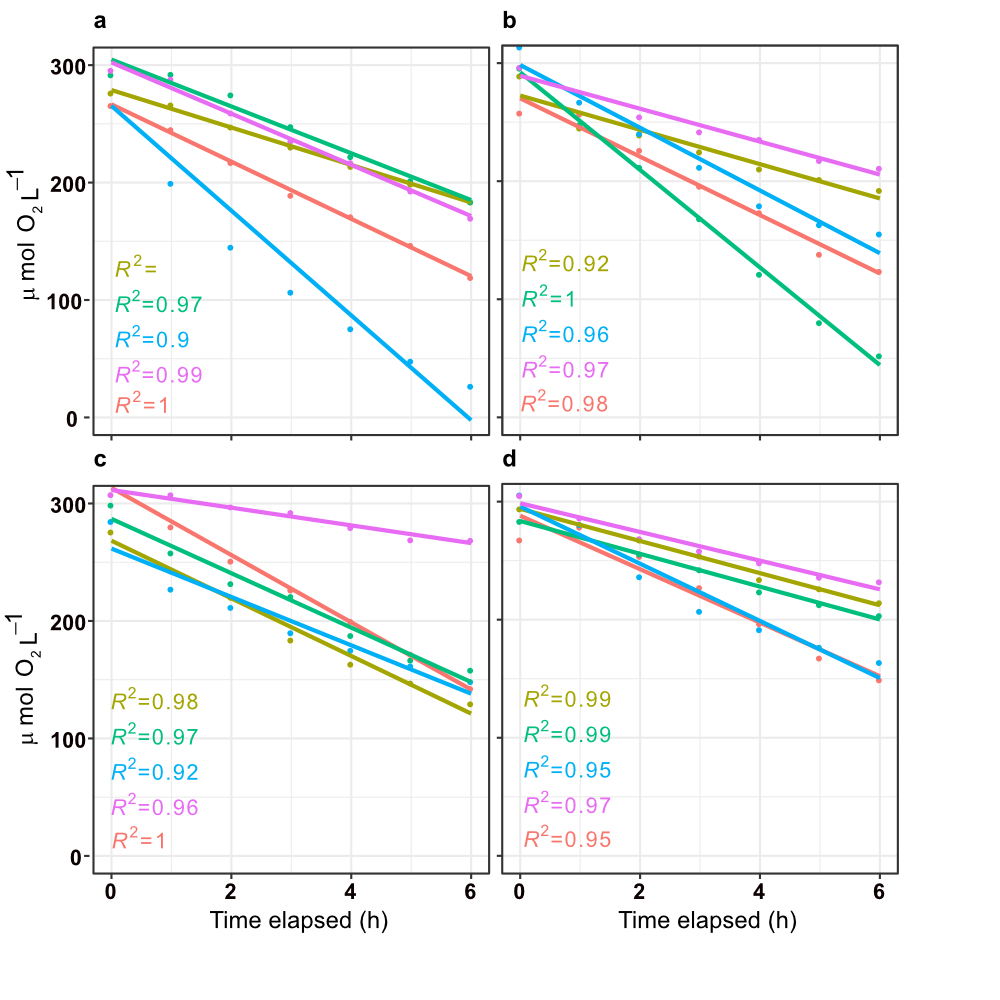

**Supplementary Fig. 4** Aerobic respiration rates (μmol Lˉ¹ hˉ¹) calculated as the slope of the linear regression between dissolved oxygen (DO) concentration and incubation time in microcosms with 0.3 mg C Lˉ¹ amendments of N-bearing compounds, (**a**) Lysine, (**b**) Serine, and N-free compounds, (**c**) Sodium Propionate, (**d**) Sodium Ascorbate. The colors in each curve are associated to a microcosm biological replicate.

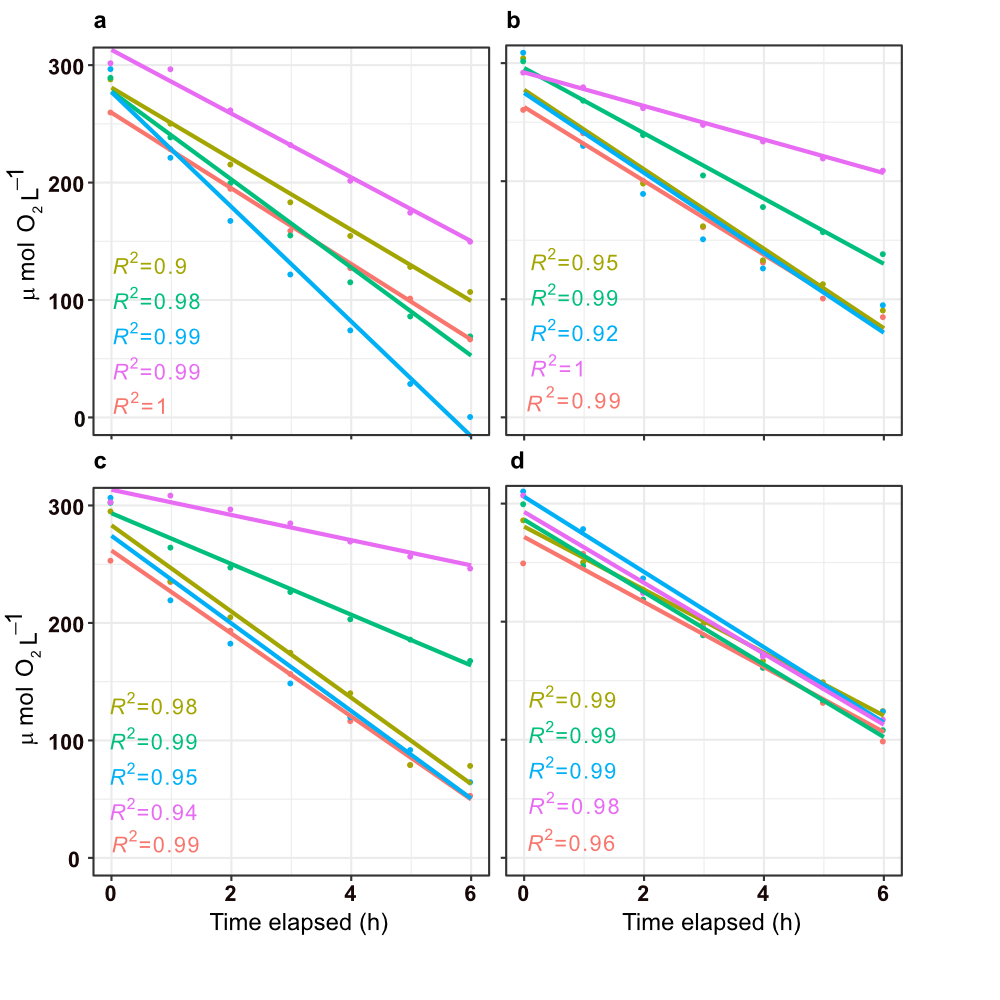
 
**Supplementary Fig. 5** Aerobic respiration rates (μmol Lˉ¹ hˉ¹) calculated as the slope of the linear regression between DO concentration and incubation time in microcosms with 3 mg C Lˉ¹ amendments of N-bearing compounds, (**a**) Lysine, (**b**) Serine, and N-free compounds, (**c**) Sodium Propionate, (**d**) Sodium Ascorbate. The colors in each curve are associated to a microcosm biological replicate.

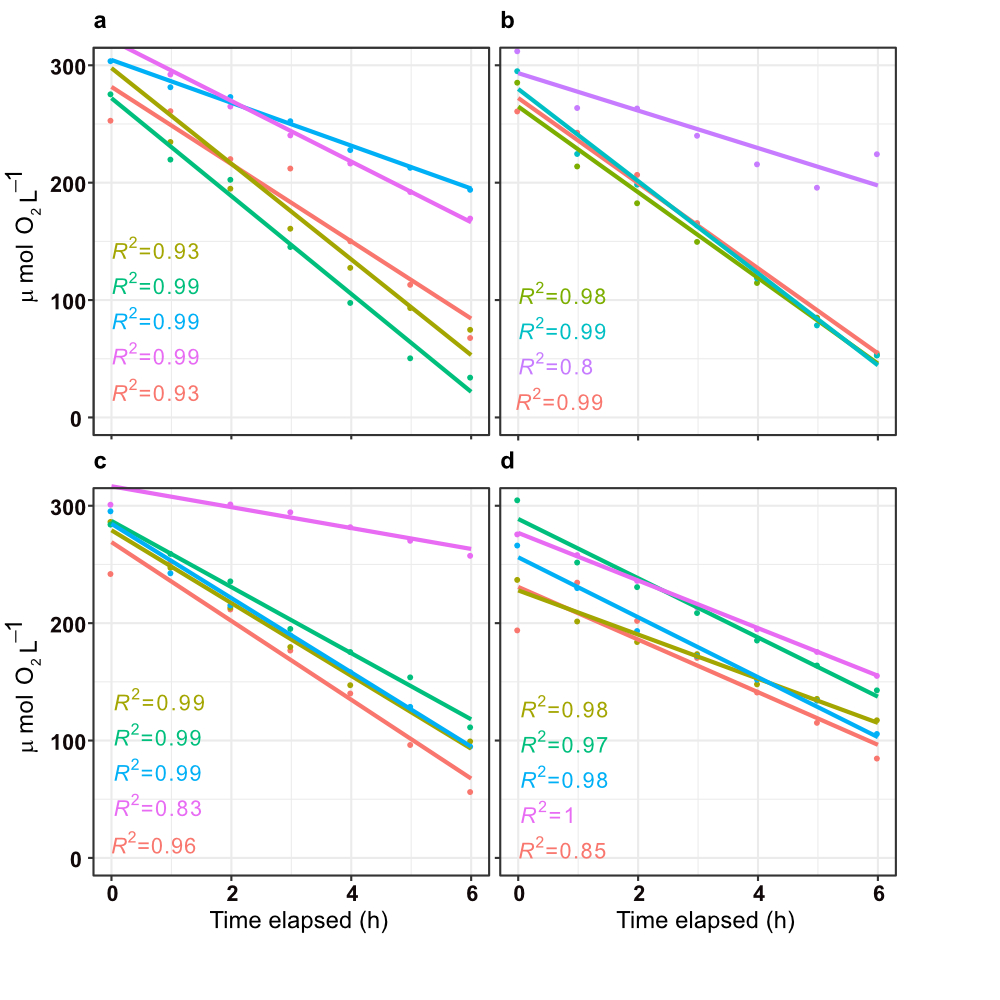

**Supplementary Fig. 6** Aerobic respiration rates (μmol Lˉ¹ hˉ¹) calculated as the slope of the linear regression between DO concentration and incubation time. Figure shows results for 9 mg C Lˉ¹ amendments of N-bearing compounds, (**a**) Lysine, (**b**) Serine, and N-free compounds, (**c**) Sodium Propionate, (**d**) Sodium Ascorbate. The colors in each curve are associated to a microcosm biological replicate.

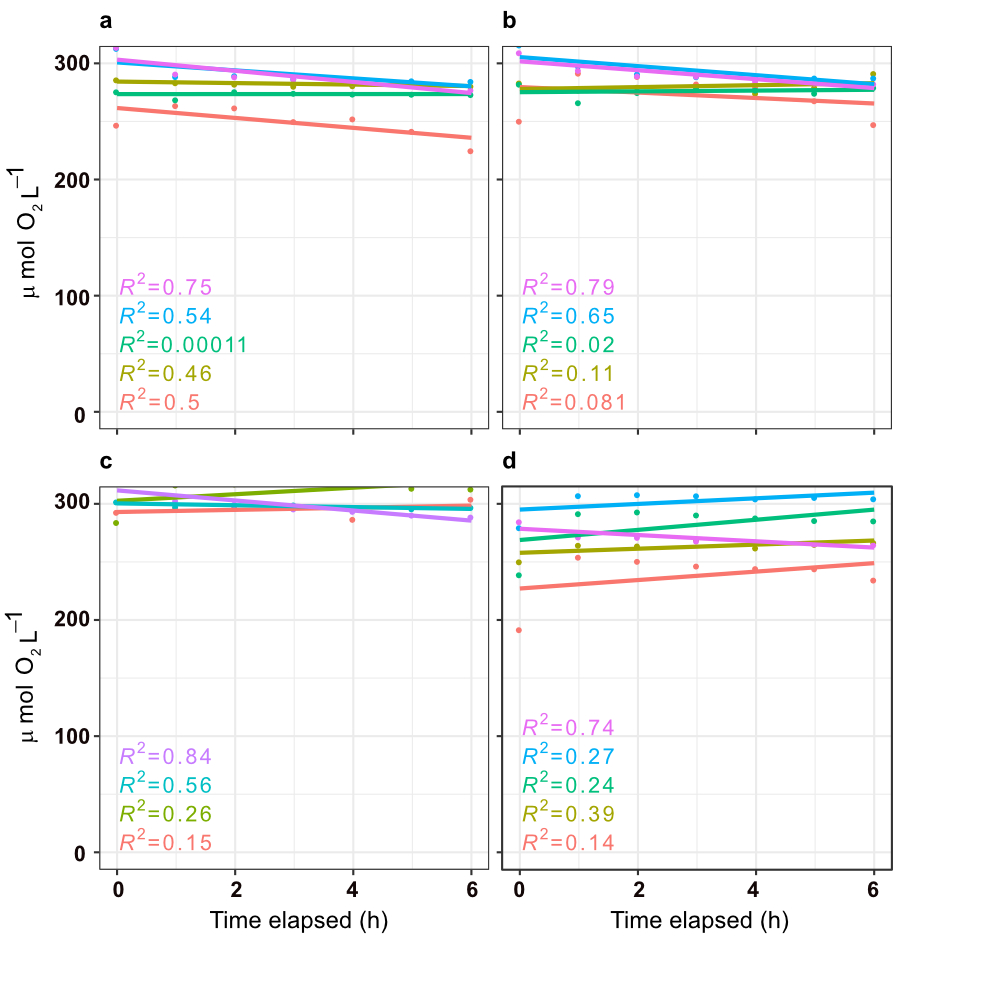

**Supplementary Fig. 7** Dissolved oxygen concentration (μmol Lˉ¹) as a function of incubation time for heat killed microcosms amended with 9 mg C Lˉ¹. The OM amendments corresponded to N-bearing compounds, (**a**) Lysine, (**b**) Serine, and N-free compounds, (**c**) Sodium Propionate, (**d**) Sodium Ascorbate. The colors in each curve are associated to a microcosm biological replicate.

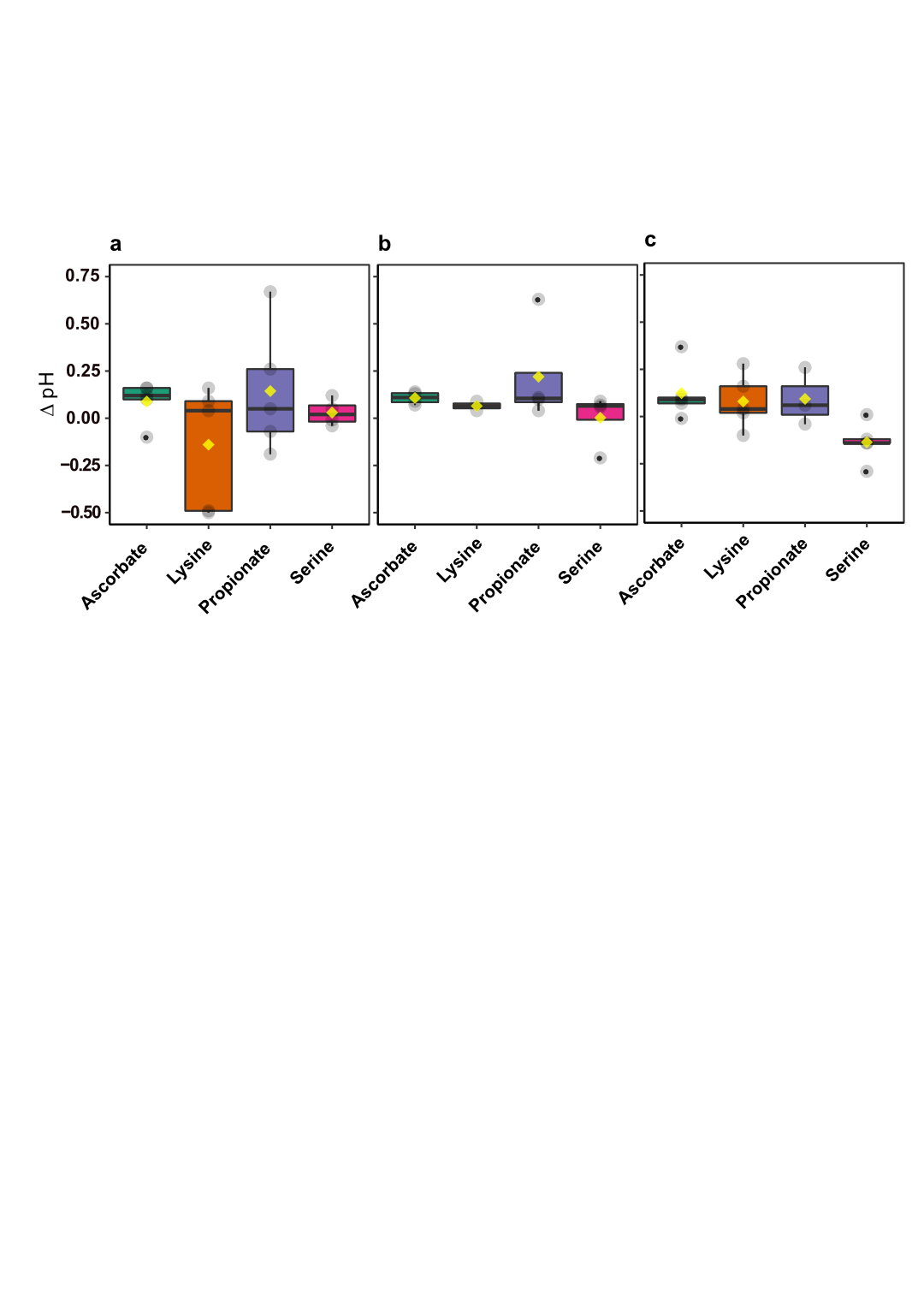

**Supplementary Fig. 8** Difference in pH (∆pH) between the beginning and end of the incubations for each OM amendment. (**a**) 0.3 mg Lˉ¹ C, (**b**) 3 mg Lˉ¹ C and (**c**) 9 mg Lˉ¹ C**.** Colors indicate different OM amendments. Plots are marked with a center black line in each box (median), yellow dot (mean), and hinges (values at 25^th^ and 75^th^ percentiles). *P*-value was derived from one-sided Mann-Whitney U test. One asterisk (*) indicates *p* ≤ 0.05, and two (**) indicates *p* ≤ 0.01.
